# An improved viability assay for *Acanthamoeba castellanii* trophozoites reveals drug-induced pseudocyst formation

**DOI:** 10.1101/2025.06.06.658351

**Authors:** Carrie A. Flynn, Rebecca I. Colón-Ríos, Andrew Harmez, Barbara I. Kazmierczak

**Affiliations:** Department of Microbial Pathogenesis, Yale School of Medicine, New Haven, CT; Department of Medicine, Yale School of Medicine, New Haven, CT

**Keywords:** free-living amoeba, *Acanthamoeba castellanii*, cell viability assays, trophozoites, cysts, pseudocysts, drug screen

## Abstract

*Acanthamoeba castellanii* is a free-living amoeba (FLA) that causes fatal human infections with few effective treatments. One limitation of new drug development is the lack of accurate, high-throughput, and quantitative viability assays that score both trophozoites and cysts. A colorimetric assay using Sulforhodamine B (SRB), which measures cell adherence, has previously been adapted for *A. castellanii* trophozoites. In this study, we demonstrated that the SRB assay can be optimized to serve as a robust, high-throughput platform that can measure viable *Acanthamoeba* trophozoites, pseudocysts and cysts. We used this assay to measure the IC50 of 18 commonly used drugs and disinfectants on *A. castellanii* trophozoites, demonstrating that several clinically used drugs induce pseudocyst formation rather than amoeba death.

## Background

*Naegleria fowleri*, *Balamuthia mandrillaris*, and *Acanthamoeba* sp. are free-living amoeba (FLA) that cause disease throughout the world. Cutaneous, ocular, disseminated, and central nervous system infections result in severe morbidity and mortality, as treatment options are limited and have high failure rates. Drugs specifically developed to treat FLA infections are critically needed (1–3), yet a remaining bottleneck in their development are assays that measure amoeba viability in a reproducible, high-throughput fashion.

Cell viability testing, which asks the seemingly simple question of whether a cell is alive or dead, is far from straightforward. Some assays measure cell viability directly (i.e. growth leading to colony or plaque formation)(4, 5). Many, however, measure cell membrane integrity(6), substrate adherence (7), or metabolic activity(8) as surrogates for viability, which in turn are affected by a cell’s physiology, differentiation state, and/or mechanism of dying(9). Senescent cells stop consuming or producing measured metabolites, as do cells that have differentiated into spores or cysts(10). Cell death can lead to membrane permeabilization or frank lysis, or can leave the membrane intact(11). Loss of adherence accompanies death, but environmental conditions and differentiation state can also cause cells to detach(12).

Measuring pathogenic FLA viability presents many challenges. FLA exist as both metabolically active, mobile trophozoites and dormant, hardy cysts. *Naegleria* species have an additional flagellate form(13), while *Acanthamoeba* spp. can rapidly respond to stress by forming pseudocysts (14). Infections with *Balamuthia* and *Acanthamoeba*, but not *Naegleria*, involve both cysts and trophozoites (15, 16). As cysts are intrinsically highly resistant to drugs, they can lead to recurrence of clinical infection despite treatment (17). Trophozoites can encyst when exposed to drugs or other stressors, exacerbating this problem(18). An optimal assay should distinguish between cysts, trophozoites, and pseudocysts and measure the viability of all forms.

Two commonly used assays, alamarBlue(19) and CellTiterGlo(20), measure *Acanthamoeba* trophozoite viability in a high-throughput manner. The only reliable way to assay cyst viability, however, is to remove the cysts from experimental conditions and demonstrate excystment by either observing trophozoites microscopically for up to a month(21) or employing a secondary trophozoite viability test (20, 22, 23). This method is slow, low-throughput, and indirect, which has motivated the search for better assays of trophozoite and cyst viability. There are no published methods for determining the viability of pseudocysts.

In this study, we tested whether an optimized sulforhodamine B (SRB) assay could serve as a robust, high-throughput platform to measure *Acanthamoeba* viability. The SRB assay measures cell adherence, potentially allowing better cyst detection than assays measuring cellular metabolism or membrane permeability. We found that this high throughput trophozoite viability assay also detected viable cells that encysted under test conditions, and discriminated live pseudocysts from dead cells. We then used this assay to determine the IC50 of 18 commonly used drugs and disinfectants on *A. castellanii* trophozoites, demonstrating the value of this assay in testing of potential amoebicides.

## Methods

### Media

Chemicals (suppliers) and media recipes are detailed in Supplemental Materials. **Culture of amoebas**. *A. castellanii* Neff strain (provided by Dr. Craig Roy, Yale University) was cultured axenically in 75 cm^2^ tissue culture-treated flasks in 40 mL PYG medium without antibiotics, without shaking at 25^◦^C. Amoeba were passaged every 3 days (1:80 dilution) for no more than 15 passages. Confluent trophozoite monolayers washed once with PYG, then harvested into 20 mL fresh PYG by tapping gently to detach cells. Cells were counted, pelleted (1,000 x *g* for 15 min) and resuspended in PYG at 10^6^ cells/mL. 96 well plates were seeded for assays with 100 µL (10^5^ cells) per well. Plates were centrifuged (200 x *g* for 5 min) and incubated at room temperature for 2 hours to allow cells to adhere to plates. All spins were done at room temperature.

Cysts were prepared by two methods. For Fig S4, confluent trophozoite monolayers in flasks were rinsed with EM, then incubated in 40 mL EM without shaking at 25^◦^C for at least 3 days until all trophozoites had encysted (as determined by microscopy). Cysts were washed, harvested by scraping in 20 mL EM and counted. Cysts were pelleted (1,500 x *g* for 20 min) and resuspended in EM at a concentration of 10^6^ cysts per mL or as needed. 96 well plates were seeded with 100 µL of cysts/well, spun (200 x *g* for 5 min), immediately fixed by adding 25 µL 50% TCA in PBS-MC to each well, and assayed by SRB.

For all other experiments, encystment was induced in 96-well plates by harvesting and plating trophozoites to 96 well plates as described above, then spinning plates (200 x *g* for 5 min) to ensure attachment. PYG medium was removed and 100 µL per well EMb was gently added gently to the walls of the wells (to avoid cell detachment). Plates were covered with breathable plate seals (Axygen BF-400 breathable sealing film) instead of lids to ensure even oxygenation and incubated at 25^◦^C until all trophozoites had encysted (>36 h, as determined by microscopic examination).

Autoclaved controls were prepared by harvesting trophozoites and cysts as above, then autoclaving cells in 10-mL glass screw-top vials for 15 minutes. Media volume lost to boiling was replaced and 100 µL per well was added to 96-well plates.

### Compound testing

96 well plates seeded with adhered trophozoites or cysts were spun (200 x *g* for 5 min); media was removed and replaced with 100 µL/well fresh media ± test compound. Plates were either fixed immediately or covered with breathable plate seals and incubated for 20 h at 25°C without shaking, as indicated.

### Optimized SRB assay

At the end of an experiment, spent media was removed from wells. (Of note, plates were not centrifuged prior to this step as it could cause detached/non-viable cells to adhere to the well bottom and generate a false positive signal.) 125 µL of 10% TCA (w/v) in PBS-MC was gently pipetted onto sides of wells and plates were incubated at 4°C for at least 1h. Plates were washed four times by submerging in large trays of tap water. Plates were dried by tapping onto paper towels. 50 µL of freshly prepared SRB dye (0.2 g SRB dissolved in 5 mL of 1% acetic acid) was added per well and incubated for 15 min at room temperature. Plates were washed 3 times by submersion in 1% acetic acid and again tapped dry on paper towels. 150 µL of 10 mM Tris-HCl, pH 8 was added per well, and plates were incubated on a gyrating platform for 5 min at room temperature to solubilize dye before reading absorbance at OD_565_.

### Pseudocyst staining

Spent media was removed from treated wells, transferred to Eppendorf tubes, and detached pseudocysts were stained with 100 µg/mL ConA-Alexa Fluor 488 for 20-30 minutes in the dark at room temperature. 5 volumes of PBS were added and samples pelleted at 20,000 x *g* for 5 min. Cell pellets were resuspended in 4 µL PBS, and 2 µL of cells spotted onto a 1.5% low-melt agarose pad, which was then placed (cell-side down) on a coverslip (24). Samples were imaged immediately using a Nikon Eclipse Ti-E inverted microscope (100x objective with oil) and a Hamamatsu ORCA-Fusion BT Digital CMOS camera using the phase-brightfield and YFP channels. (If imaging is delayed, any live trophozoites will migrate to the edges of the agar pads where there is more oxygen and leave the field of view.)

### Statistics and data analysis

Data was graphed and analyzed using GraphPad Prism version 10. Mean and standard error of the mean (SEM) are displayed for each scatterplot. Statistical significance was determined for two groups using a two-tailed, unpaired t test with Welch’s correction and for three or more groups using one-way Brown-Forsythe and Welch ANOVA tests (for unequal SDs) either Dunnett’s T3 (for n<50 per group) or Games-Howell (for n>50 per group) comparisons tests; an alpha of 0.05 was used to determine significance.

## Results

### Optimization of the sulforhodamine B (SRB) assay for high throughput detection of viable *A. castellanii*

The sulforhodamine B (SRB) assay was originally developed for use on cancer cells(7). The assay was later applied to *A. castellanii* trophozoites(25) to measure live, adherent cells in a 96-well format(25–27) (Fig S1). This simple assay, in which trichloroacetic acid-fixed cell proteins are stained with the fuchsia aminoxanthene dye SRB under weakly acid conditions, is inexpensive, quick to perform, high-throughput (7) and robust to environmental conditions (pH, media composition) and cell metabolism (25). The three major published SRB protocols differ significantly (Table S1)(7, 25, 26), and have not been tested on cysts. We first sought to optimize this assay for trophozoites and then assess its utility for measuring cyst viability. Optimization is described in Supplemental Results (Fig S2 -S7) and led to the optimized SRB assay used in subsequent experiments.

### Optimized protocol allows for accurate determination of number of amoebas

The SRB assay was used to determine viable amoebae number, generating standard curves for live trophozoites (Fig 1, blue) and cysts (Fig 1, red). Absorbance correlated well with the number of live amoebas, with R^2^ values >0.99 for both trophozoites and cysts. The lower limit of detection for the assay was 1,000 amoebas, allowing for accurate detection of >99% killing when starting with 100,000 cells per well.

**Figure 1:**
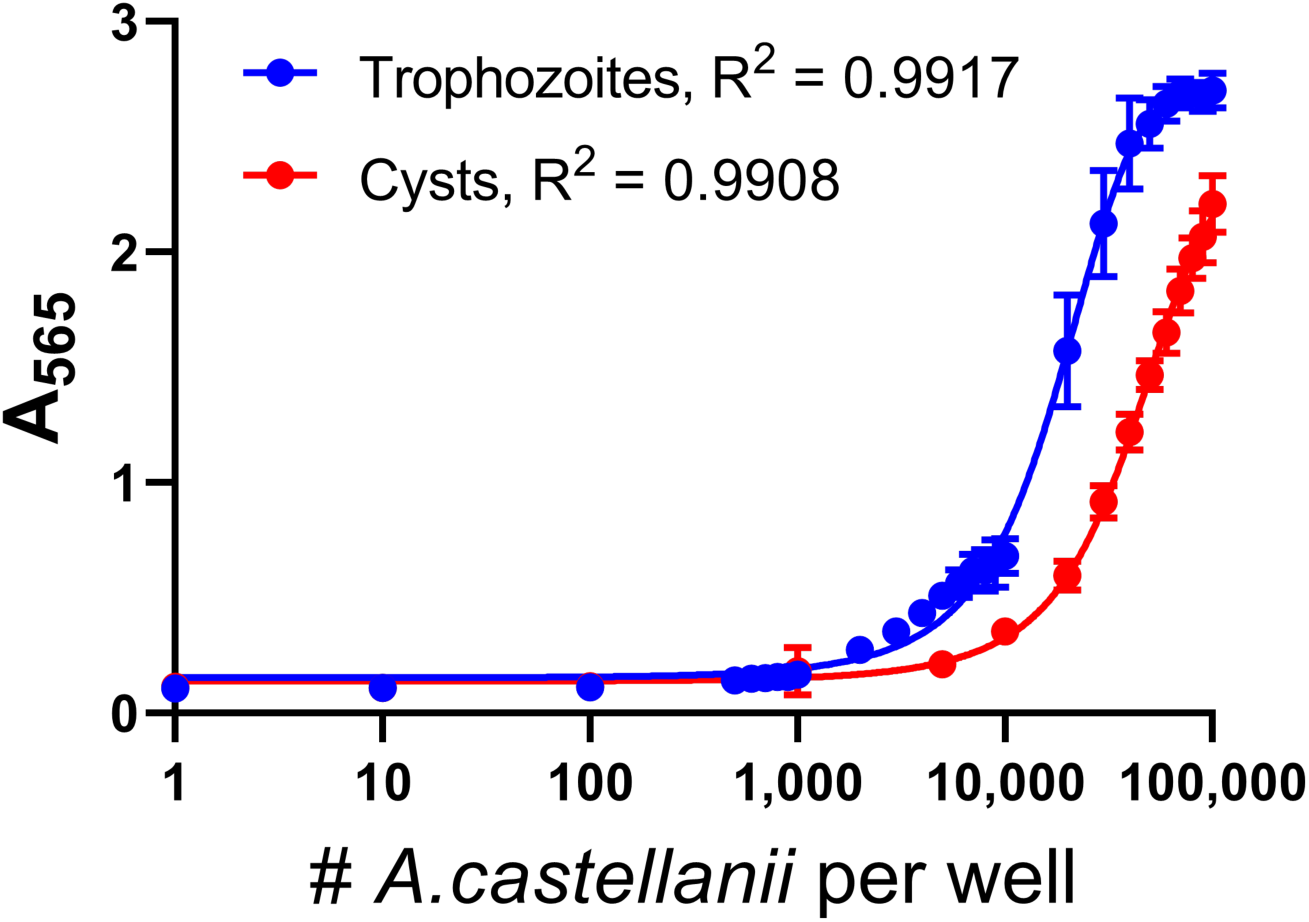
The optimized SRB assay has excellent sensitivity for detecting live trophozoites and cysts. Dilution curve of trophozoites (blue) and cysts (red) using the final optimized protocol (media removed before adding 10% TCA in PBS-MC, 50 µL per well 4% SRB, submerged washes). Data represent the mean +/-S.D. of at least 4 independent experiments with at least 4 replicates each. Nonlinear regression was performed (Gompertz growth curve with least squares fit); R^2^ values displayed on graph.

Although cysts and trophozoites stained similarly with SRB (Fig S4D), absorbance readings consistently differed when comparing the same number of trophozoites plated to a well in PYG vs. seeded to a well and encysted in situ (Fig 1). We ascribed this difference to cell loss during encystment and estimated that our protocol yielded approximately 50,000 cysts for every 10^5^ trophozoites initially seeded to the well.

### The SRB assay underestimates cyst death

The ability of the SRB assay to discriminate between live vs. dead cysts was compared to the gold standard, i.e. outgrowth assays (Fig 2). Amoeba encysted in duplicate 96 well plates were treated with drugs or disinfectants and assayed in parallel by SRB assay and outgrowth. For the latter, media plus non-attached cells were removed from wells and separately cultured from residual (attached) cells to reveal the presence of viable cysts in each compartment after treatment. We included live cyst controls as well as cysts killed by various methods to assess whether the SRB assay could measure different types of cell death.

**Figure 2:**
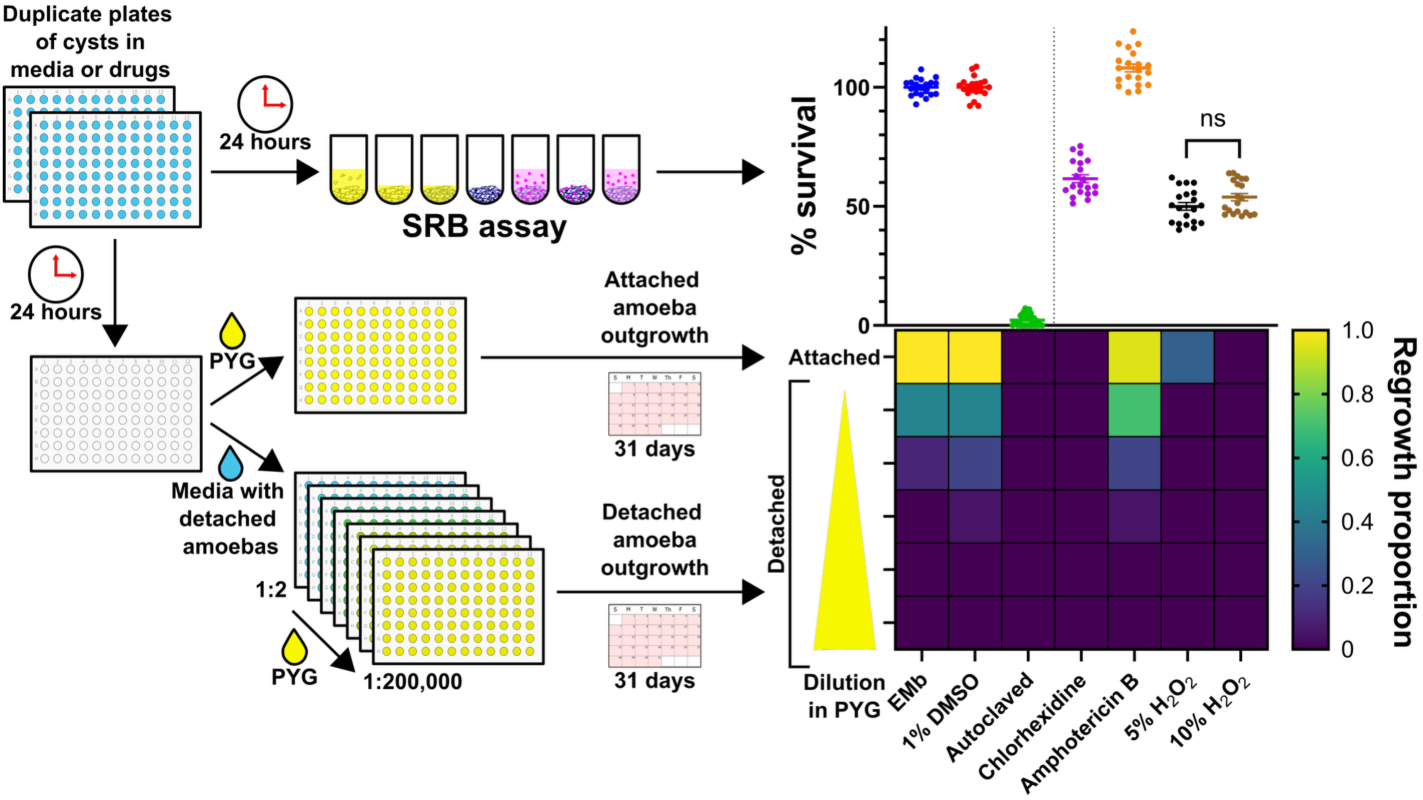
Outgrowth experiments reveal underestimation of cyst death by SRB assay. Schematic of experiment illustrates experimental design and corresponding results. 10^5^ trophozoites per well were encysted in EMb in 96-well plates for 30 hours at 25^◦^C, as described above. Duplicate plates of cysts were incubated for 24 hours at 25^◦^C in conditions as indicated. For one plate, spent media was removed, cells were fixed, and the SRB assay was performed (scatter plot). For the second plate, the spent media was transferred to a new plate, mixed 1:1 with fresh PYG, and serially diluted. Fresh PYG was also added back to wells of the original plate (attached). Plates were incubated and monitored for growth for 31 days, with fresh PYG added daily to feed cells and maintain volume. The proportion of wells that regrew during the month was scored for each condition (heat map). Amphotericin B (lipid nanosphere formulation, “LNS-AmB”) and chlorhexidine were dissolved in DMSO and added at 1% final volume in EMb (200 µM amphotericin B, 100 µM chlorhexidine); H_2_O_2_ was diluted in EMb. Scatter plot reports percent survival measured by SRB assay. Data expressed as the percentage of cells surviving at the end of incubation compared to the mean A565 value for the condition-appropriate control wells (EMb or EMb + 1% DMSO) for each experiment. The mean and SEM of at least 3 independent experiments with at least 4 replicates per condition are shown. Statistical comparisons were calculated using a one-way ANOVA with Tukey’s correction for multiple comparisons in GraphPad Prism software; ns = not significant. Heat map reports outgrowth of viable amoebas by showing the proportion of wells that regrew during the month-long incubation.

The SRB assay reported high viability for media control wells (EMb with or without 1% DMSO) and low viability for autoclaved cysts (Fig 2, scatter plot). Expected regrowth of amoebas was observed for both attached and detached compartments in media and vehicle controls (Fig 2, heat map). Wells seeded with autoclaved cysts showed no outgrowth after 31 days. Amphotericin B (200 µM) showed no cysticidal activity, with full survival scored by SRB assay (Fig 2, scatter plot) and outgrowth of both attached and detached compartments (Fig 2, heat map) similar to media controls. A greater proportion of wells regrew in the original plate for each condition than for the replated media, consistent with larger numbers of live amoebas adhering to wells than removed with the media.

Although there was little (5% H_2_O_2_) or no (10% H_2_O_2_ and chlorhexidine) regrowth of cysts treated with disinfectants (Fig 2, heat map), the SRB assay scored ∼50% survival of cysts treated with these agents (Fig 2, scatter plot). This discrepancy–a high signal from the SRB assay despite confirmation of cyst death–was likely due to adherence of proteinaceous outer cyst wall material that stained strongly with SRB dye despite cyst killing (Fig S8)(28, 29). Thus, the SRB assay could not measure cysticidal activity. However, the SRB assay did accurately detect live cysts, and thus could correctly score trophozoites encysting in response to treatment as live instead of dead.

### Outgrowth experiments confirmed that the SRB assay measures live, not dead trophozoites

SRB assay performance for *A. castellanii* trophozoites was also evaluated by performing outgrowth experiments on cells treated in duplicate with chemical disinfectants or drugs (Fig 3). Media ± 1%DMSO and autoclaved trophozoites served as positive and negative controls. Cell morphology was examined by microscopy at the end of incubation, before removing spent media.

**Figure 3:**
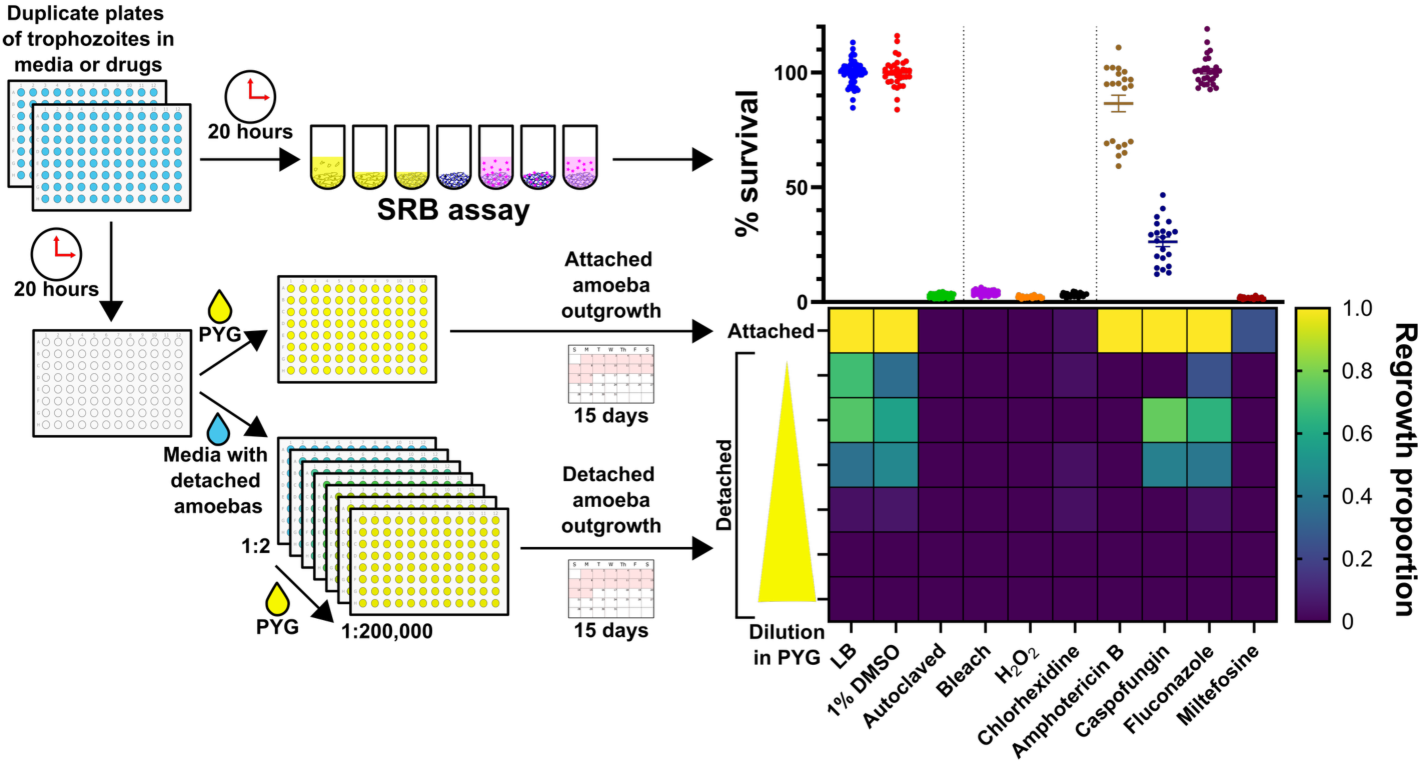
**The SRB assay measures live, not dead, trophozoites**. Schematic of experiment illustrates experimental design and corresponding results. 10^5^ trophozoites per well were incubated in duplicate 96-well plates for 20 hours at 25^◦^C in conditions as indicated. For one plate, spent media was removed, cells were fixed, and the SRB assay was performed (scatter plot). For the second plate, the top 50 µL of spent media was transferred to a new plate and the remainder was removed and discarded. The transferred media was mixed 1:1 with fresh PYG, and serially diluted. Fresh PYG was also added back to wells of the original plate (attached). Plates were incubated and monitored for growth for 15 days, with fresh PYG added daily to feed cells and maintain volume. The proportion of wells that regrew during this time was scored for each condition (heat map). Amphotericin B (LNS-AmB), fluconazole, miltefosine, chlorhexidine, and caspofungin were dissolved in DMSO; bleach and H_2_O_2_ were dissolved in PBS; all were added at 1% final volume in LB at the following concentrations: 1% bleach, 10% H_2_O_2_, 20 µM chlorhexidine, 256 µM amphotericin B, 64 µM caspofungin, 512 µM fluconazole, 256 µM miltefosine. **Scatter plot:** Percent survival measured by SRB assay. Data expressed as the percent of cells surviving at the end of incubation compared to the mean A565 value for the condition-appropriate control wells (LB or LB + 1% DMSO) for each experiment. The mean and SEM of at least 3 independent experiments with 7 replicates per condition are shown. **Heat map:** Outgrowth of viable amoebas. Heat map shows the proportion of wells that regrew during the 15-day incubation.

The SRB assay reported high survival for media control wells and low survival for autoclaved trophozoites (Fig 3, scatter plot). Bleach and H_2_O_2_ demonstrated robust killing of trophozoites, leaving dead cells with a granular appearance and non-intact membranes (Fig S9). Of the tested drugs, only chlorhexidine and miltefosine showed strong trophocidal activity by SRB assay, supported by microscopic visualization of dead cells. Amphotericin B and fluconazole appeared inactive, while caspofungin showed moderate killing. By microscopy, cells treated with fluconazole were indistinguishable from media controls. Most cells incubated in amphotericin B were live trophozoites, though a minority of cells developed a rounded, non-adherent morphology.

Neither the wells containing autoclaved trophozoites in the original plate nor their removed and replated media showed any outgrowth after 15 days, as expected (Fig 3, heat map). No growth in the original plate or of replated media at any dilution was detected for bleach or H_2_O_2_, confirming the SRB assay finding of full killing (Fig 3, scatter plot). All other conditions yielded outgrowth, indicating the presence of live cells that were either attached (if in the original plate) or non-adherent (if in the removed, serially diluted and replated media). As with cysts, there was more regrowth of attached than detached trophozoites for each condition, consistent with a greater number of live amoebas adherent to the wells than removed with the spent media.

Fluconazole-treated trophozoites showed outgrowth of both attached and detached cells like that seen for the LB and 1% DMSO media controls (Fig 3, heat map), consistent with the high survival reported by SRB assay (Fig 3, scatter plot). Regrowth of attached cells occurred for all amphotericin B wells, demonstrating resistance of trophozoites to this drug (consistent with SRB assay results (Fig 3, scatter plot)).

A small proportion of wells treated with miltefosine (25%) regrew, indicating that a small number of live, attached cells remained after treatment. Chlorhexidine was highly amoebicidal, with only 1 of 28 replicate wells exhibiting growth of adhered cells, consistent with the high activity reported by SRB assay.

Caspofungin was the only condition for which results of the SRB assay and the outgrowth experiment showed significant disagreement. Although the SRB assay reported 26% mean survival, outgrowth of both adherent and media/detached compartments was similar to media controls. This result suggested that caspofungin might induce formation of pseudocysts, a non-adherent *Acanthamoeba* life form that would not be detected as “viable” by the SRB assay but which would show outgrowth upon replating.

In summary, outgrowth experiments confirmed that SRB absorbance was strongly correlated with the presence of viable amoebas. Second, media removed prior to fixation had few viable cells, such that loss of adherence was a reliable proxy for trophozoite death. Lastly, comparison of SRB and outgrowth assay results suggested that some drug treatments might induce pseudocyst formation.

### A method to detect *A. castellanii* pseudocysts

To determine whether some drugs caused pseudocyst formation, we added a step to our method that distinguished live pseudocysts from dead cells, relying on selective staining of pseudocyst membranes by fluorescent concanavalin A (ConA). This lectin binds the two major components coating the pseudocyst envelope, mannose and glucose (14). Live trophozoites were unstained or took up ConA exclusively in their vacuoles, while autoclaved cells stained diffusely with ConA (Fig S10). Staining of pseudocysts (induced by exposure to 3% DMSO for 2 hours and confirmed by phase imaging(14)) was restricted to the membrane surface, creating a distinctive round fluorescent halo.

Caspofungin (64 µM) treated samples showed mixtures of trophozoites, pseudocysts, and dead cells following ConA straining, as suspected from the results of SRB and outgrowth experiments (Fig 4). Staining of LNS-AmB (256 µM) treated cells showed that most were live trophozoites, though some samples contained pseudocysts clustered with trophozoites. We wondered whether LNS-AmB induced membrane alterations that made pseudocysts adhere to trophozoites, thereby preventing most from being removed with spent media that was replated to detect outgrowth of detached cells (Fig 3). Other drugs did not cause appreciable pseudocyst formation at these high concentrations (Fig 4).

**Figure 4:**
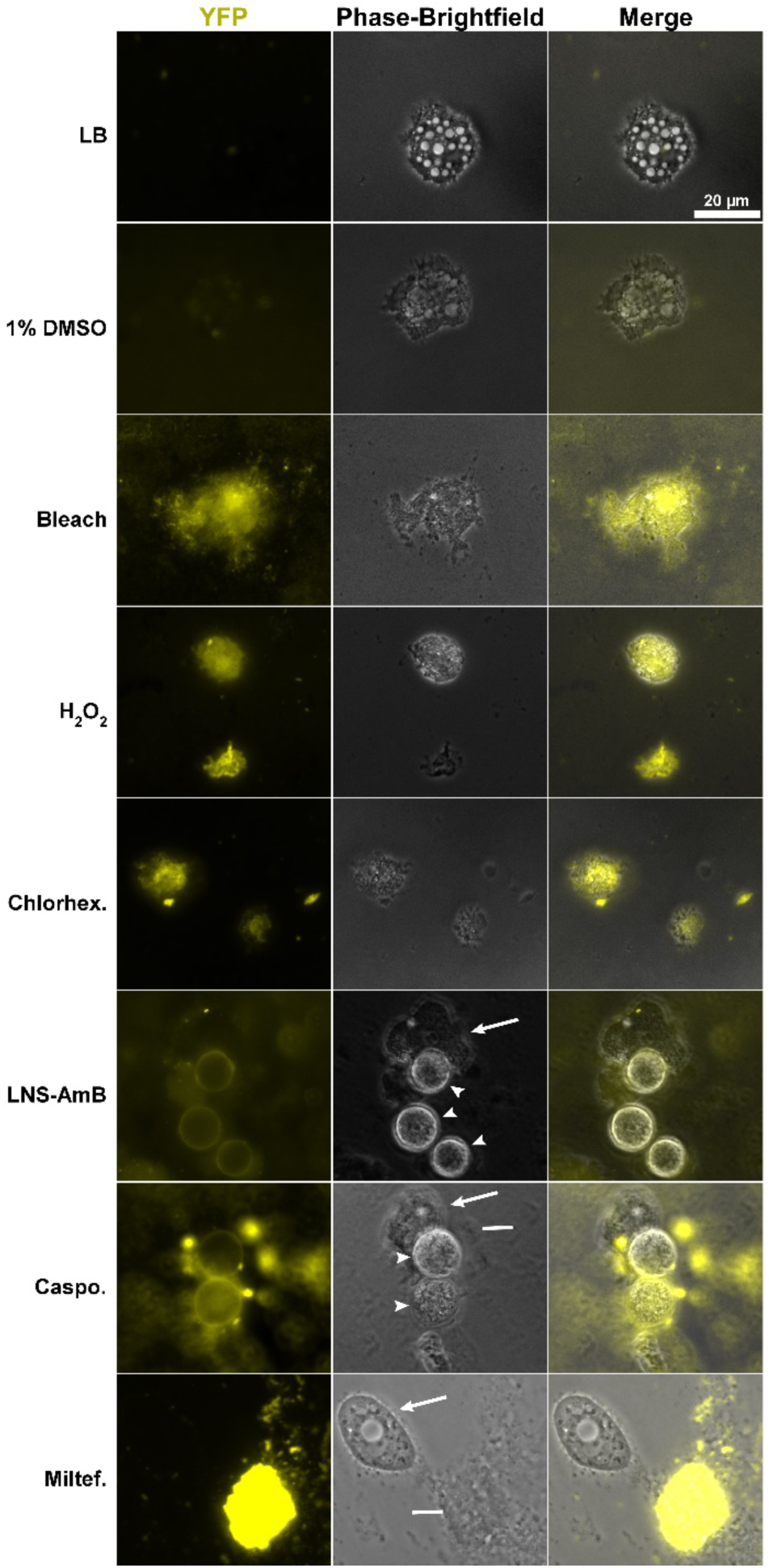
Fluorescent ConA staining reveals pseudocyst induction by caspofungin and amphotericin. **B.** 10^5^ trophozoites per well were incubated in a 96-well plate for 20 hours at 25^◦^C in conditions as indicated. Non-adherent cells in spent media were stained with ConA-Alexa Fluor 488 and imaged using a Nikon Eclipse Ti-E inverted microscope (100x objective with oil) and a Hamamatsu ORCA-Fusion BT Digital CMOS camera. Amphotericin B, miltefosine, chlorhexidine, and caspofungin were dissolved in DMSO; bleach and H_2_O_2_ were diluted with PBS; all were added at 1% final volume in LB to achieve the following concentrations: 1% bleach, 10% H_2_O_2_, 20 µM chlorhexidine, 256 µM amphotericin B, 64 µM caspofungin, 256 µM miltefosine. Representative images displayed. Arrows, live trophozoites; arrowheads, live pseudocysts; lines, dead cells.

### Applying the SRB assay to measure trophocidal drug activity

Commercially available antibiotic, antiparasitic, and antifungal agents with demonstrated or hypothesized activity against *A. castellanii* were tested on trophozoites at a range of concentrations, as were chemical disinfectants with reported activity (Fig 5). IC50s were calculated for each drug that killed at least 50% of the cells and compared to published values (Table 1). Phase contrast microscopy was used to confirm morphological phenotypes of live and dead cells (Fig S11), while detached cells were stained with ConA to detect pseudocysts (Fig S12).

**Figure 5:**
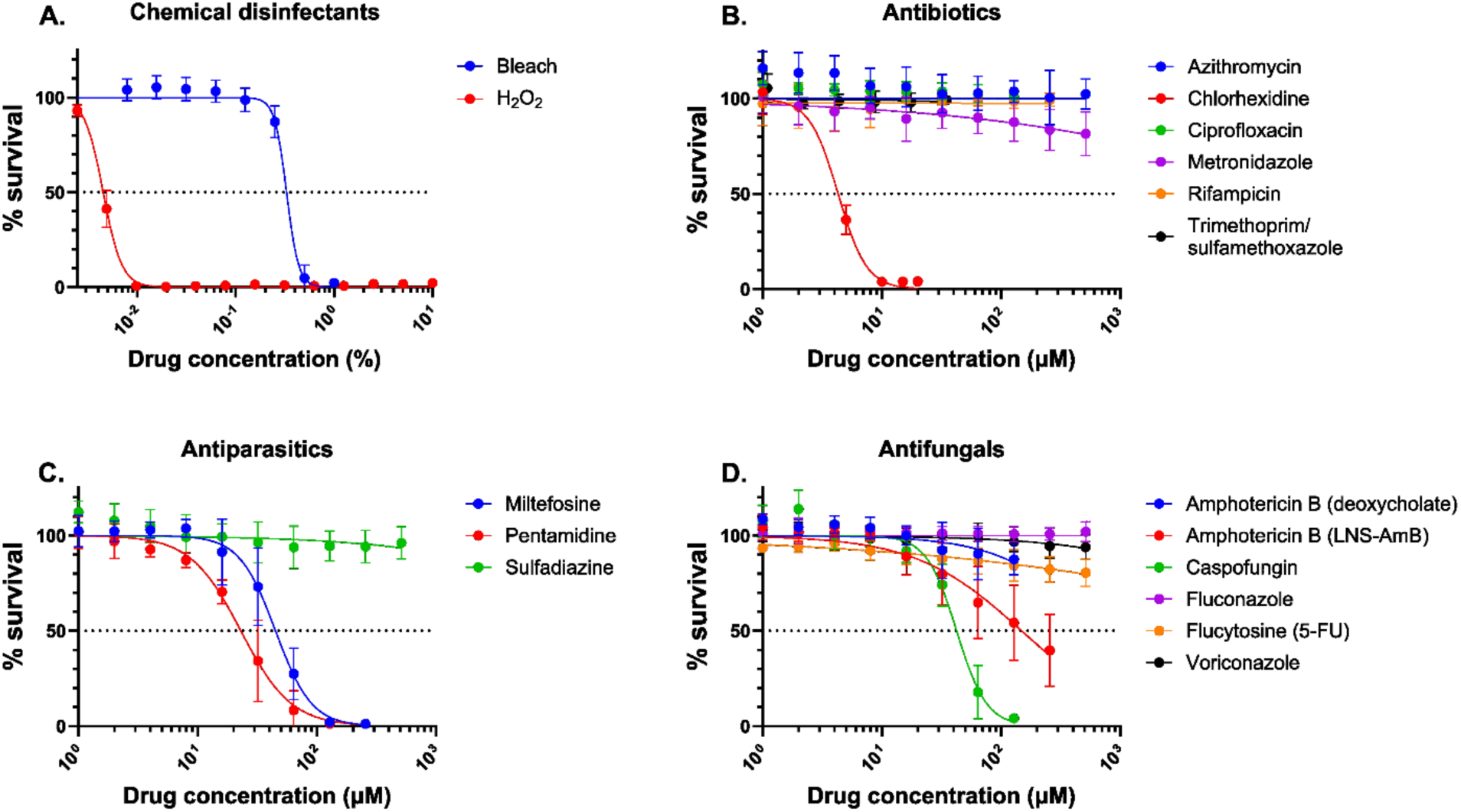
**Assessing the trophocidal activity of drugs using the SRB assay**. 10^5^ trophozoites per well were incubated in 96-well plates for 20 hours at 25^◦^C in conditions as indicated. At the end of the incubation, spent media was removed and discarded, cells were fixed, and the SRB assay was performed. Amphotericin B (lipid nanosphere formulation, “LNS-AmB”), fluconazole, voriconazole, miltefosine, chlorhexidine, sulfadiazine, azithromycin, ciprofloxacin, metronidazole, rifampicin, trimethoprim/sulfamethoxazole, and caspofungin were dissolved in DMSO; bleach, H_2_O_2_, pentamidine, and flucytosine were dissolved in PBS; amphotericin B (deoxycholate formulation) was purchased as a solution in water. Two-fold dilutions were prepared, and drugs were added at 1% final volume in LB at a range of concentrations (listed in Table 1) according to their solubility and activity. Trimethoprim was combined with sulfamethoxazole at a ratio of 1:5.7; concentration plotted is of trimethoprim. Data expressed as the percent of cells surviving at the end of incubation compared to the mean A565 value for the condition-appropriate control wells (LB or LB + 1% DMSO) for each experiment. The mean and SD of at least 3 independent experiments with at least 3 replicates per condition are shown.

**Table 1.**
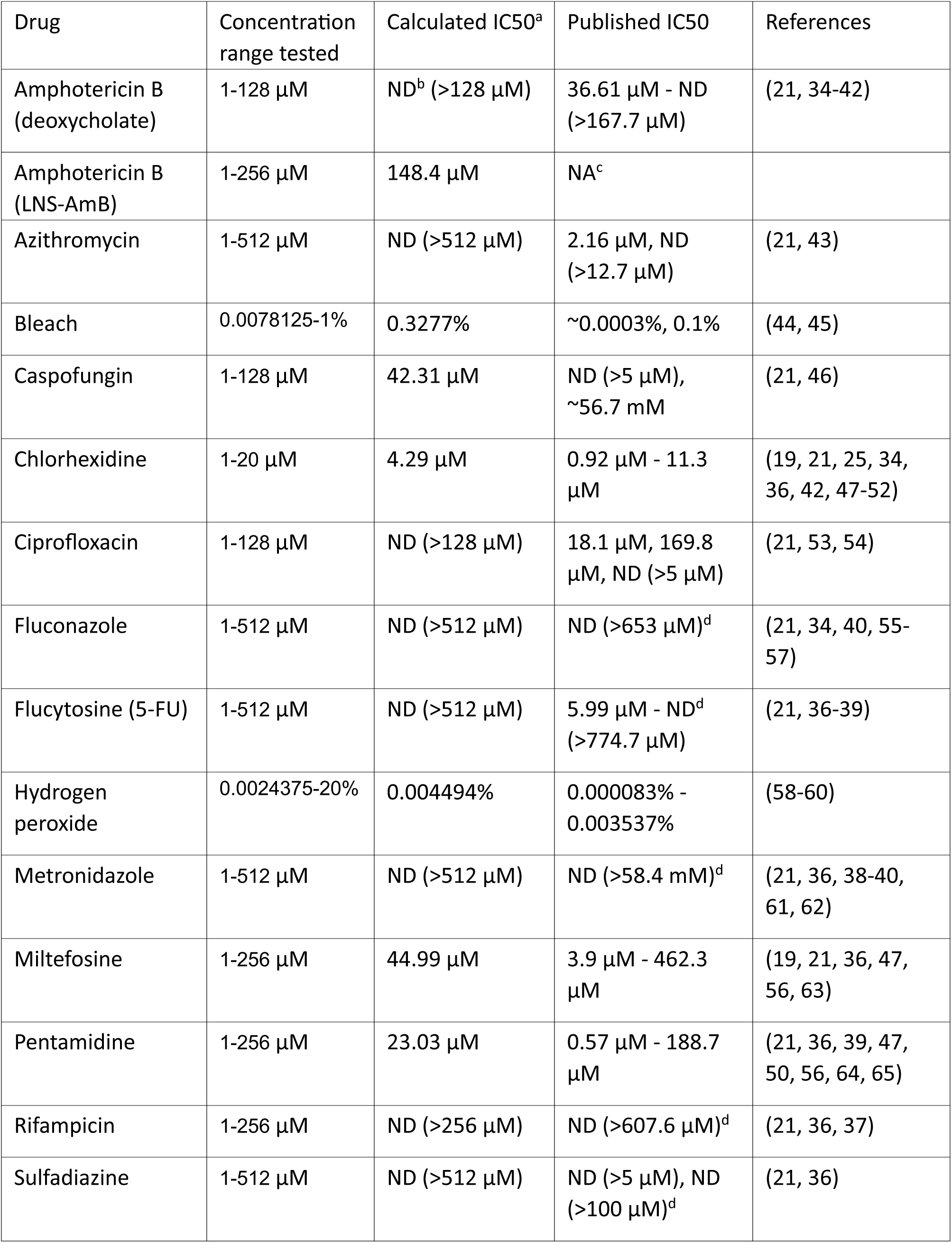
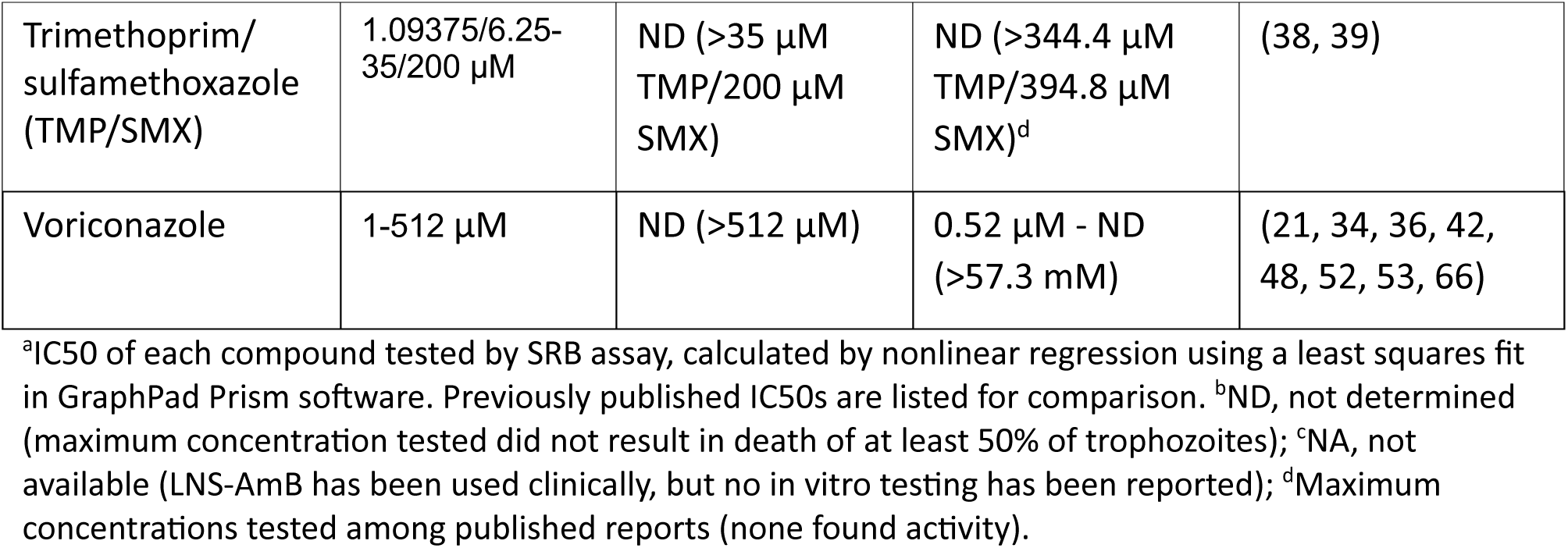
IC50 of tested compounds.

| Drug | Concentration range tested | Calculated IC50<sup>a</sup> | Published IC50 | References |
| --- | --- | --- | --- | --- |
| Amphotericin B (deoxycholate) | 1-128 μM | ND<sup>b</sup> (>128 μM) | 36.61 μM - ND (>167.7 μM) | (21, 34-42) |
| Amphotericin B (LNS-AmB) | 1-256 μM | 148.4 μM | NA<sup>c</sup> | |
| Azithromycin | 1-512 μM | ND (>512 μM) | 2.16 μM, ND (>12.7 μM) | (21, 43) |
| Bleach | 0.0078125-1% | 0.3277% | ~0.0003%, 0.1% | (44, 45) |
| Caspofungin | 1-128 μM | 42.31 μM | ND (>5 μM), ~56.7 mM | (21, 46) |
| Chlorhexidine | 1-20 μM | 4.29 μM | 0.92 μM - 11.3 μM | (19, 21, 25, 34, 36, 42, 47-52) |
| Ciprofloxacin | 1-128 μM | ND (>128 μM) | 18.1 μM, 169.8 μM, ND (>5 μM) | (21, 53, 54) |
| Fluconazole | 1-512 μM | ND (>512 μM) | ND (>653 μM)<sup>d</sup> | (21, 34, 40, 55-57) |
| Flucytosine (5-FU) | 1-512 μM | ND (>512 μM) | 5.99 μM - ND<sup>d</sup> (>774.7 μM) | (21, 36-39) |
| Hydrogen peroxide | 0.0024375-20% | 0.004494% | 0.000083% - 0.003537% | (58-60) |
| Metronidazole | 1-512 μM | ND (>512 μM) | ND (>58.4 mM)<sup>d</sup> | (21, 36, 38-40, 61, 62) |
| Miltefosine | 1-256 μM | 44.99 μM | 3.9 μM - 462.3 μM | (19, 21, 36, 47, 56, 63) |
| Pentamidine | 1-256 μM | 23.03 μM | 0.57 μM - 188.7 μM | (21, 36, 39, 47, 50, 56, 64, 65) |
| Rifampicin | 1-256 μM | ND (>256 μM) | ND (>607.6 μM)<sup>d</sup> | (21, 36, 37) |
| Sulfadiazine | 1-512 μM | ND (>512 μM) | ND (>5 μM), ND (>100 μM)<sup>d</sup> | (21, 36) |
| Trimethoprim/<br>sulfamethoxazole<br>(TMP/SMX) | 1.09375/6.25-35/200 μM | ND (>35 μM<br>TMP/200 μM<br>SMX) | ND (>344.4 μM<br>TMP/394.8 μM<br>SMX)<sup>d</sup> | (38, 39) |
| Voriconazole | 1-512 μM | ND (>512 μM) | 0.52 μM - ND<br>(>57.3 mM) | (21, 34, 36, 42,<br>48, 52, 53, 66) |

<sup>a</sup>IC<sub>50</sub> of each compound tested by SRB assay, calculated by nonlinear regression using a least squares fit in GraphPad Prism software. Previously published IC<sub>50</sub>s are listed for comparison. <sup>b</sup>ND, not determined (maximum concentration tested did not result in death of at least 50% of trophozoites); <sup>c</sup>NA, not available (LNS-AmB has been used clinically, but no in vitro testing has been reported); <sup>d</sup>Maximum concentrations tested among published reports (none found activity).

Kill curves demonstrated that both bleach and H_2_O_2_ were active against trophozoites at concentrations slightly higher than those previously reported (Fig 5A, Table 1). Of the seven antibiotics tested (Fig 5B, Table 1), only chlorhexidine had activity, with an IC50 within the range of published reports. Microscopy confirmed that all three of these active agents resulted in dead cells at doses corresponding to the drop in SRB absorbance value (Fig S11). In agreement with the literature (Table 1), we detected no trophocidal activity for metronidazole, rifampicin, or trimethoprim/sulfamethoxazole. Neither azithromycin nor ciprofloxacin had trophocidal activity by SRB assay; the literature contains conflicting reports of activity for these agents (Table 1). For all these inactive antibiotics, monolayers of live trophozoites were visualized by microscopy at all tested concentrations (Fig S11).

The antiparasitics miltefosine and pentamidine demonstrated trophocidal activity, while sulfadiazine did not (Fig 5C). The IC50 values of these active drugs fell within the broad ranges previously reported (Table 1). Miltefosine appeared to induce pseudocysts at concentrations of 32-128 µM (when SRB absorbance values declined) before killing all cells at 256 µM (Fig S12).

Two of the six antifungal drugs tested (Fig 5D) were trophocidal, with moderate (caspofungin) or weak activity (LNS-AmB). Caspofungin’s IC50 (as determined by SRB assay) was ∼1000-fold lower than that previously reported, while results of in vitro LNS-AmB testing have not been reported to our knowledge (Table 1). Both caspofungin and LNS-AmB treatment resulted in significant pseudocyst formation (Fig S11, S12). The traditional deoxycholate formulation of amphotericin B, widely reported to lack activity against *Acanthamoeba*, was also inactive by our assay. Both tested azoles (fluconazole and voriconazole) lacked activity; this agrees with published reports for fluconazole, but reports for voriconazole vary widely. Similarly, flucytosine was inactive in our hands, but published testing shows a wide range of IC50 values (Table 1).

Overall, only seven of the 18 drugs and chemical disinfectants tested demonstrated trophocidal activity and only one - chlorhexidine - was highly active with an IC50 less than 10 μM.

## Discussion

Determination of amoeba viability is technically difficult(22): a method must account for trophozoites, pseudocysts, and cysts despite their distinct biologic and metabolic properties. The optimized SRB assay, coupled with a step to detect pseudocysts, reliably measures live trophozoites, pseudocysts, and cysts and accounts for dead trophozoites and pseudocysts via a rapid and high throughput method that is also inexpensive and highly reproducible. The SRB assay is compatible with bleach or H_2_O_2_, common positive controls for killing, and accurately scores cells killed by various mechanisms (autoclaving, chemical disinfectants, and drugs), as confirmed by outgrowth experiments, microscopy, and targeted membrane staining.

Using the SRB assay, we tested 18 potential amoebicides and calculated IC50s for active agents. Bleach, H_2_O_2_, chlorhexidine, and pentamidine efficiently killed trophozoites, with IC50 values similar to or higher than those reported by other groups. This variation could be attributable to differences in amoeba strain, assay method, and/or testing conditions (which are not standardized). Our method, however, avoided falsely scoring either live cysts or pseudocysts as dead, a testing error that would result in a lower apparent IC50.

Three agents—LNS-AmB, miltefosine, and caspofungin—induced pseudocyst formation which we detected with ConA staining (Fig 4, Fig S12). Increasing concentrations of LNS-AmB resulted in increasing numbers of pseudocysts, few dead cells, and many trophozoites; however replated media showed no outgrowth despite the presence of pseudocysts (Fig 3). These findings suggest that LNS-AmB may cause cell surface changes that promote cell-cell adherence, giving rise to the clusters of pseudocysts and trophozoites we observed and limiting pseudocyst detachment. SRB and outgrowth assays report the presence of these viable, drug-resistant cells (Fig 3). Miltefosine induced many pseudocysts at intermediate concentrations (32-128 µM); at 256 µM, most cells were dead though a small population of viable trophozoites persisted (Fig S12), consistent with SRB and outgrowth assay results (Fig 3). These observations suggest that susceptible cells form pseudocysts before dying in response to miltefosine, while a subpopulation of intrinsically resistant cells remain trophozoites. Treatment with caspofungin likewise induced pseudocysts at intermediate doses (64 µM), while a higher dose (128 µM) of drug showed many dead cells, rare pseudocysts and no viable trophozoites. SRB and outgrowth assays conducted at this intermediate dose were discordant (Fig 3) – with decreased viability by SRB but significant outgrowth of viable cells – consistent with the presence of viable, non-adherent pseudocysts on ConA staining (Fig S12).

The scoring of cell adherence by the SRB assay as a proxy for viability led to false positive results when cells detached without dying, e.g. during pseudocyst formation (as detailed above). We overcame this limitation by adding an additional fluorescent staining step, which accurately identified live trophozoites, live pseudocysts, and dead cells. The assay yielded false negative results when attached cysts were killed, due to staining of proteinaceous exocyst material by the SRB dye. To our knowledge, this inability to account for live vs. dead cysts is a drawback our assay shares with other published methods (save outgrowth-based assays). We therefore advise against using the optimized SRB method as a cyst viability assay and recommend using incubation times that are too short to allow complete encystment (such as the 20-hour duration we selected in this study). This assay, however, will not mistakenly score encysting cells as “dead” and is therefore likely to produce fewer false positive hits during drug screens on *A. castellanii* trophozoites than other methods.

This study provides a new option—high throughput, reproducible, and rigorously characterized—for viability testing of *A. castellanii* trophozoites. It also highlights the persistent need to develop new amoebicidal drugs, as no systemic agent killed *A. castellanii* at doses achievable in humans(30). Our results also question the utility of several standard therapeutic agents (fluconazole, sulfadiazine, and flucytosine), as well as other drugs (i.e. voriconazole, trimethoprim/sulfamethoxazole, macrolides) reportedly used against systemic *Acanthamoeba* infections. While other *Acanthamoeba* strains may be sensitive, the Neff strain is fully resistant to these drugs; additional testing using this pipeline on other species and strains is warranted. Finally, our drug activity testing demonstrated that three agents induce pseudocyst formation. While others have reported pseudocyst induction by novel compounds or contact lens disinfecting solutions(31–33), we believe this is the first report of pseudocyst formation stimulated by these clinically important drugs.

## Notes

### Competing Interest Statement

The authors have declared no competing interest.

