## Supplemental Materials for "An improved viability assay for *Acanthamoeba castellanii* trophozoites reveals drug-induced pseudocyst formation"

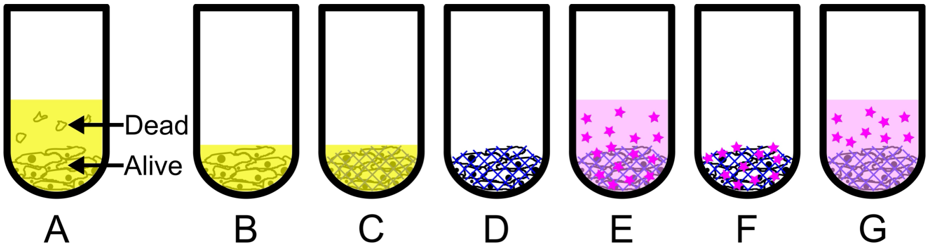


**Figure S1: The SRB assay measures amoeba viability by measuring adhered cells.** A) An individual microtiter plate well showing live, adherent *A. castellanii* cells and dead, floating cells. B) Media containing nonadherent cells is removed. C) Adherent cells are fixed to the well with trichloroacetic acid (TCA). D) TCA is washed away with water. E) Fixed amoebas are incubated in SRB dye, which binds proteins nonspecifically. F) Excess SRB dye is removed with 1% acetic acid. G) SRB dye bound to amoeba proteins is solubilized with Tris-HCL pH 8.0, and absorbance is measured.

| Reference | [7] | [26] | [25] |
|---|---|---|---|
| Cell type | >60 human tumor cell lines | BCA-1, KB, MCF-7, and Vero cell lines | *A. castellanii* (Neff T4 and clinical AK T3 strains) |
| Plate type | 96-well | 96-well | 96-well |
| Cell density (#cells per well) | 1,000-200,000 | 19,000 | 50-5,000 |
| Media removed before fixing? | No | No | For chlorhexidine test only |
| Final TCA concentration | 10% | 3.3% | 10% |
| Fixation | 1 hour at 4°C | 1 hour at 4°C | 1 hour 4°C |
| Water wash | 5x with tap water | 4x with slow-running tap water | 3x with slow-running tap water |
| SRB dye (w/v in 1% acetic acid) | 0.2-1.6% | 0.057% | 4% |
| SRB volume (µL per well) | 32-128 | 100 | 25 |
| SRB incubation time | 2.5-40 minutes | 30 minutes | 15 minutes |
| 1% acetic acid wash | 1-7x (pour into wells from beaker, flick plate) | 4x (“as quickly as possible”) | 3x (unclear method) |

| | | | |
|---|---|---|---|
| Dye solubilization | 10 mM unbuffered Tris base (pH 10.5) | 10 mM unbuffered Tris base (pH 10.5) | 10 mM unbuffered Tris base (pH 10.5) |
| Tris volume (µL per well) | Unknown | 200 | 150 |
| Tris incubation | 5 minutes on gyratory shaker | 5 minutes on gyratory shaker | 5-10 minutes on gyratory shaker |
| Absorbance reading | 564 nm | 510 nm | "dual wavelength mode": 562 nm test/630 nm reference |
| Sensitivity (#cells per well) | 2,500-200,000 | 1,000-2,000 | 100-2,000[a] |

**Table S1: Comparison of Published SRB Assay Protocols.** [a]Note: the authors propose that 500 cells per well is the optimal seeding density at which the trophozoites are confluent, but in our experience nearly $10^5$ trophozoites are required for confluence in the wells of a 96-well plate.

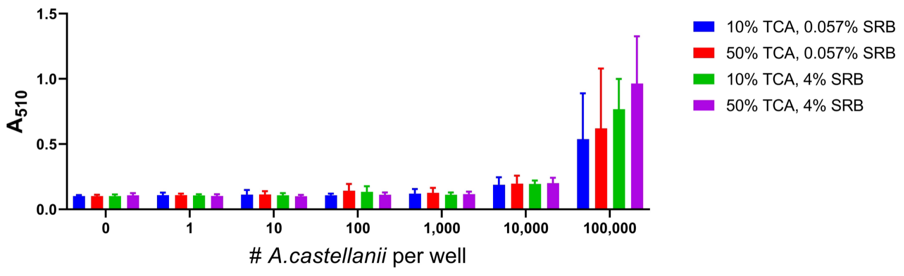


**Figure S2: Higher concentrations of TCA and SRB result in higher absorbance readings of live trophozoites.** 0-$10^5$ live trophozoites in PYG were fixed by adding 10% or 50% TCA without removing the media first (yielding final TCA concentrations of 2% and 10%, respectively). The SRB assay was performed using 25 µL SRB per well (0.057% or 4%). Plates were washed with water by holding them sideways under gently running tap water, and with acetic acid using a serological pipette and flicking off the liquid into a sink. Absorbance was measured at 510 nm. Data expressed as the mean +/- S.D. of 3 independent experiments with 3 replicates per condition.

The concentrations of TCA and SRB used in prior publications(7, 25, 26) were tested as indicated. A two-way ANOVA reveals that neither TCA nor SRB concentration is a significant contributor to variation between groups. This is likely due to high variability in the absorbance reading within each group, which we hypothesized was due to nonuniform exposure to acetic acid during the wash step. We selected 50% TCA and 4% SRB for subsequent experiments because this condition had the highest absorbance readings at the upper limit of cell density, providing for increased sensitivity over a larger range of amoebas per well.

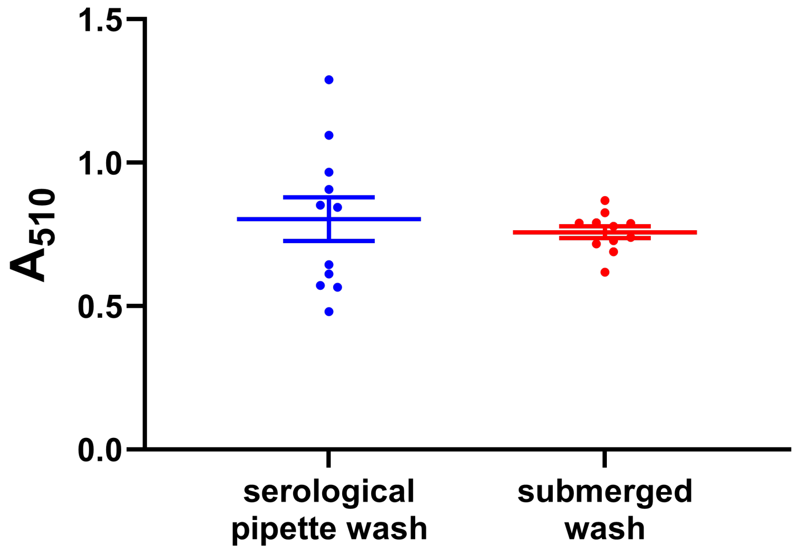


**Figure S3: Improved reproducibility by submerging plates to wash.** $10^5$ live trophozoites per well were incubated for 24 hours at 25°C in LB and fixed by adding 25 µL per well 50% TCA directly to wells without removing the media first. The SRB assay was performed using 25 µL per well of 4% SRB. Plates were washed using a serological pipette or by submerging in shallow 2000-mL plastic trays. Data are expressed as mean and SEM of 4 independent experiments with 3 replicates per condition.

Previously published methods(7, 25, 26) advise rinsing plates with water under a gently running tap, though they note that a direct jet of water to the bottom of the well can detach the cell monolayer. Additionally, they highlight the importance of washing with acetic acid as quickly as possible to avoid overbleaching, but do not outline the best way to do this. We initially tried using a multichannel pipette, but this was slow and cumbersome. A serological pipette did not perform much better and introduced too much variation in volume and exposure time between wells resulting in heterogeneous absorbance values (Fig S2). We tested a different way of washing plates by submerging in shallow plastic trays, using this method for both the water and acetic acid washes (Fig S3). We found this new method of washing was the fastest and most uniform way to wash plates (Video S1). In addition to improved speed and ease of plate processing, washing the plates in this manner decreased heterogeneity between wells, enhancing the reproducibility of the assay.

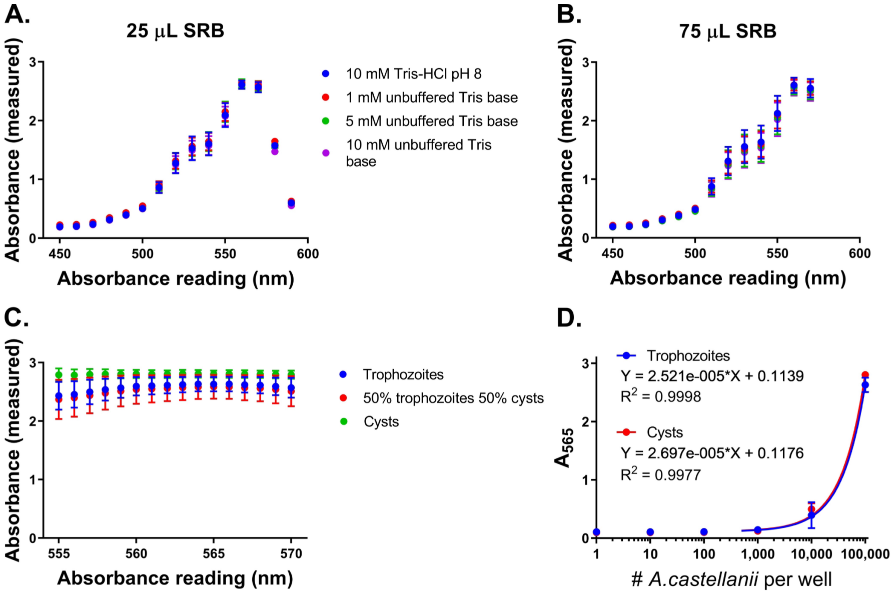


**Figure S4: Optimizing absorbance reading, Tris concentration, and SRB volume yields an accurate, reproducible assay for live trophozoites and cysts.** A, B) $10^5$ live trophozoites per well were added to plates and fixed by adding 25 µL per well 50% TCA directly to wells without removing the media first. The SRB assay was performed using 25 µL (A) or 75 µL (B) of 4% SRB dye and the indicated concentrations of Tris-HCl or unbuffered Tris base. Absorbance was measured from 450-590 nm at intervals of 10 nm. Data represent the mean +/- S.D. of 3 independent experiments with at least 4 replicates per condition. C) $10^5$ live cells (trophozoites, cysts, or equal parts both) per well were added to plates and fixed by adding 25 µL per well 50% TCA directly to wells without removing the media first. The SRB assay was performed using 50 µL SRB dye and 10 mM Tris-HCl. Absorbance was measured from 555-570 nm at intervals of 1 nm. Data represent the mean +/- S.D. of 3 independent experiments with at least 8 replicates per condition. D) Dilution curve of 0-$10^5$ live trophozoites or cysts per well, fixed by adding 25 µL per well 50% TCA directly to wells without removing the media first, using the optimized SRB assay parameters (50 µL per well 4% SRB dye, submerged washes, 150 µL per well 10 mM Tris-HCl, absorbance at 565 nm). Data represent the mean +/- S.D. of 3 independent experiments with at least 8 replicates per condition. Linear regression was performed; equations of best fit lines and $R^2$ values are displayed on the graphs.

The optimal absorbance wavelength, Tris concentration, and volume of SRB dye were then determined. For experiments performed using 25 or 75 µL SRB dye, there was no difference between the four concentrations of Tris tested in the linear range of absorbance readings, as determined by a one-way ANOVA with Tukey's multiple comparisons test (Fig S4A,B). We therefore decided to perform our experiments using 10 mM Tris-HCl pH 8 for ease of use and to reduce variability between batch preparations of Tris solution.

We then compared the two volumes of SRB dye for the 10 mM Tris-HCl data points at the range of optimal absorbance readings (550, 560, and 570 nm). An unpaired, two-tailed t-test at each of the three absorbances revealed no significant difference between the two volumes (Fig S4A,B). We therefore selected the minimum volume needed to easily cover the entire bottom of the well, 50 µL.

After determining the maximum absorbance readings to be in the range of 550-570 nm by performing an absorbance scan at 10 nm intervals (Fig S4A,B), we repeated the experiment (using the SRB dye volume and Tris-HCl concentration described above) in this narrower window at 1 nm intervals (Fig S4C) for trophozoites, cysts, and a mixture of the two. Because a compound may induce encystation or excystation, it is important that the assay measures the two cell types equally. The measured absorbances are most similar between the three groups (trophozoites, cysts, and mixed) when read at 565 nm, as determined by a two-way ANOVA with Tukey's multiple comparisons test (Fig S4C). We then performed a linear regression on the measured absorbance values for a concentration gradient of trophozoites and cysts at 565 nm, which demonstrates excellent fit (Fig S4D). An unpaired, two-tailed t-test comparing the two cell types revealed no significant difference between them. Therefore, the SRB assay is able to accurately measure live A. castellanii cells in both the trophozoite and cyst forms.

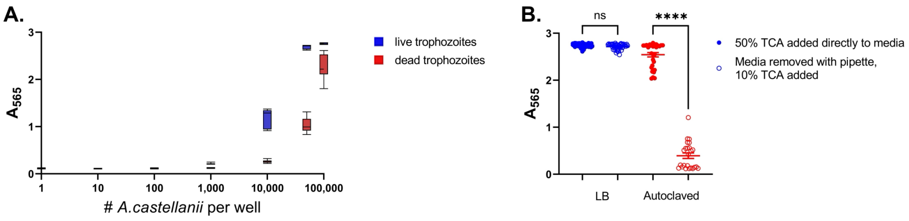


**Figure S5: Removing media before fixing with TCA prevents measurement of dead trophozoites.** A) Dilution curve of 0-$10^5$ live or autoclaved (dead) trophozoites per well in PYG medium. Cells were fixed immediately by directly adding TCA to wells without removing media first. Box and whiskers plot represents the median and 5th-95th percentiles of 2 independent experiments with at least 4 replicates per condition. B) $10^5$ live or autoclaved (dead) trophozoites were incubated for 24 hours in LB medium and fixed by either adding 50% TCA directly to wells without removing the media first (10% final TCA concentration, closed circles) or removing the spent media before adding 10% TCA in LB (open circles). Data represent the mean and SEM of 3 independent experiments with at least 5 replicates per condition. Statistical differences calculated with one-way ANOVA with Tukey's correction for multiple comparisons; ns for not significant ($p > 0.05$), **** for $p < 0.0001$.

We next tested the performance of the optimized assay on cells killed by autoclaving. Dead trophozoites were aliquoted to 96-well plates, then fixed by the addition of 25 µL of 50% TCA per well(7, 25, 27). Unexpectedly, A565 values were only slightly lower for dead trophozoites than for live trophozoites (Fig S5A). We reasoned that dead cells that settled on the bottom of the well could be fixed with TCA, even if they were not adherent, and give a false reading with the SRB assay. We further modified the assay to remove media (and non-adherent cells) before fixing with TCA. This modification did not change the A565 values for live cells, but significantly decreased the absorbance measured for dead (autoclaved) amoebae when compared to leaving media in place (Fig S5B). This supported our hypothesis that floating cells are removed with the media while live, adhered cells remain attached. We therefore incorporated the removal of media and non-adherent cells from wells before fixation.

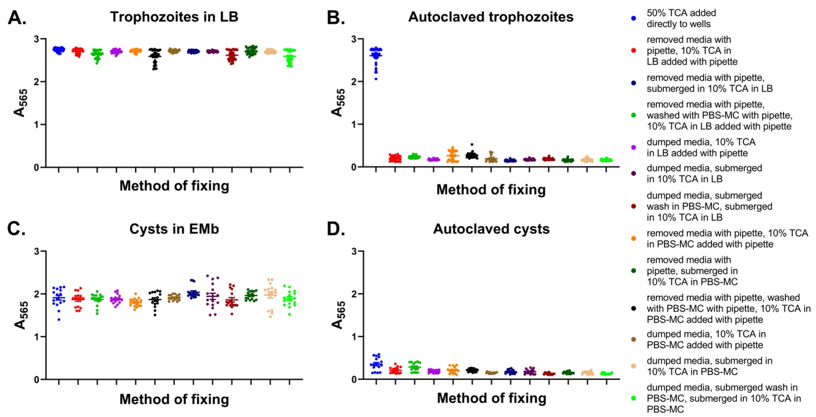


**Figure S6: Removing media by any mechanism before fixing with TCA prevents measurement of dead trophozoites and cysts.** $10^5$ live or dead (autoclaved) trophozoites or cysts per well were incubated for 24 hours at 25°C in LB (trophozoites) or EMb (cysts). Cells were fixed as indicated and the SRB assay was performed. Data are expressed as the mean and SEM of at least 3 experiments with at least 4 replicates per condition.

We next compared 13 different methods of fixing live and dead trophozoites and cysts. Amoebas killed by autoclaving were compared with live trophozoites or cysts in their standard experimental media, LB or EMb, respectively. We found that all 13 methods resulted in similar absorbance readings for live trophozoites and cysts (Fig S6A,C). We again observed that adding TCA without first removing the media resulted in elevated absorbance readings for both dead trophozoites and cysts (Fig S6B,D). The remaining twelve methods, all of which included media removal before fixation, worked equally well. Varying how media was removed (by aspiration or flicking off), adding an additional wash step, carrying out TCA fixation in rich media (LB) or buffer (PBS-MC), or adding TCA to wells more or less vigorously had no further effect on SRB assay performance, but did allow for faster processing of plates. This underscores the importance of removing the floating, dead cells before adding TCA to the wells.

Removing media before fixation provided additional advantages. If TCA was added directly to the media, plates could not be stored at 4◦C before processing as floating dead cells could settle and become fixed to the wells over time, resulting in falsely elevated absorbance measurements(7). Also, media removal prior to fixation avoided interference by proteinaceous media components such as serum(7).

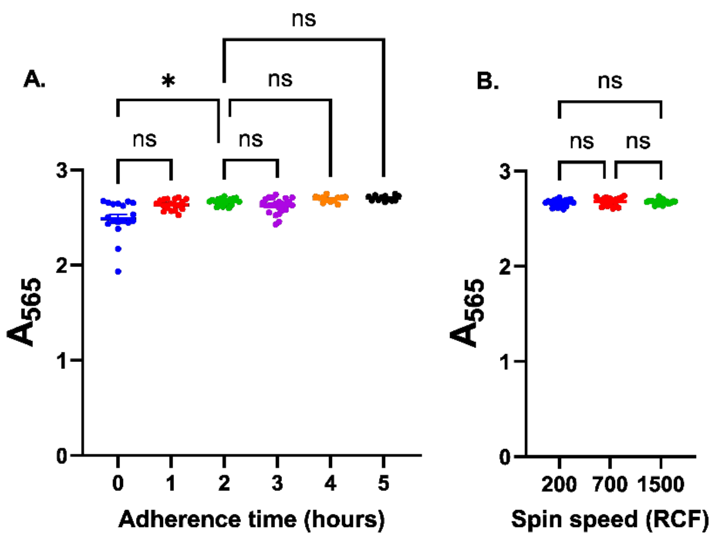


**Figure S7: Trophozoites require 2 hours to adhere to plates. A:** Trophozoites were added to plates and spun at 200 xg for 5 minutes before incubating at room temperature for 0-5 hours; media was then removed, and cells were fixed. **B:** Trophozoites were added to plates and spun at 200, 700, or 1500 xg before incubating at room temperature for 2 hours, removing media, and fixing cells. **A, B:** Scatter plots report the absorbance measured by SRB assay. The mean and SEM of 3 independent experiments with 6 replicate wells per condition are shown. Statistical differences calculated with one-way ANOVA (Brown-Forsythe and Welch ANOVA tests for unequal SDs) with Dunnett's T3 multiple comparisons test; ns for not significant ($p>0.05$), * for $p<0.05$.

Because the SRB assay relies on cell adherence, we reasoned that cells should ideally make contact with the well bottoms soon after plating and may require time to express adhesion proteins and properly attach to the wells. We therefore tested different centrifugation speeds and adherence times--during which plates were incubated undisturbed at room temperature before changing the media--to optimize trophozoite attachment (Fig S7).

After centrifuging plates containing trophozoites at 200, 700, or 1500 RCF, there was a significantly greater number of adherent cells if they were given 2 or more hours to attach before changing the media (Fig S7A; graph shows results for plates spun at 200 xg, but the same pattern was observed at all spin speeds). The speed of centrifugation was not significant if cells were then given 2 hours to attach (Fig S7B). We therefore selected a centrifugation speed of 200 xg, using the minimum necessary speed to avoid stressing cells unnecessarily, and an attachment time of at least 2 hours before changing the media for all future experiments.

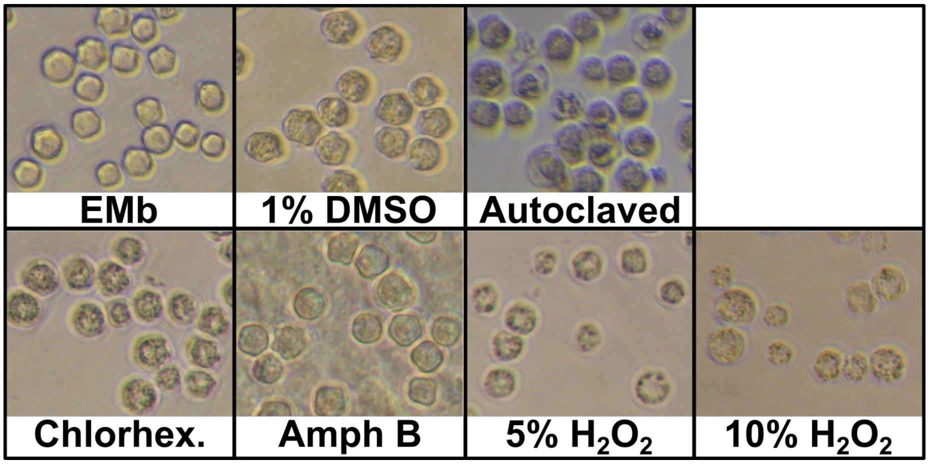


**Figure S8: The ectocyst remains upon cell death.** Microscopy of cysts in original plate after 24-hour incubation with drugs or media, as described in figure 2. Images acquired before spent media was removed from wells using a Canon Vixia HFS200 camera and Nikon Eclipse TS100 microscope (20x objective). Precipitated amphotericin B is visible deposited on the plate in the background of image (Amph B).

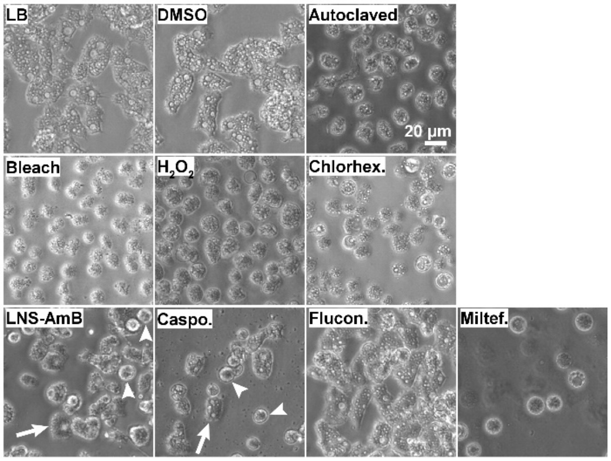


**Figure S9: Microscopy confirms phenotypes of live and dead cells after drug treatment.** Microscopy of trophozoites in original plate after 20-hour incubation with drugs or media, as described in figure 3. Phase images acquired before spent media was removed from wells using an EVOS FL Auto Imaging System (20x objective). Caspofungin well shows a small amount of precipitated drug in the background of image. Arrows, live trophozoites; arrowheads, live pseudocysts.

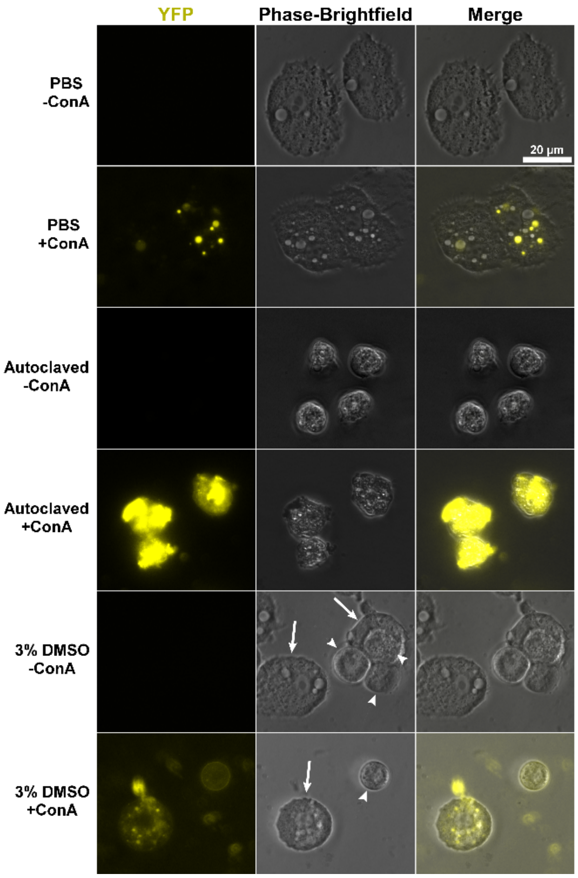


**Figure S10: Fluorescent ConA staining discriminates pseudocysts from dead cells.** Trophozoites were autoclaved or incubated in PBS or PBS + 3% DMSO for 2 hours (a duration too short for encystment), stained with 100 µg/mL ConA-Alexa Fluor 488 (+ConA) or PBS vehicle (-ConA), and imaged using a Nikon Eclipse Ti-E inverted microscope (100x objective with oil) and a Hamamatsu ORCA-Fusion BT Digital CMOS camera in a MatTek glass-bottom dish overlaid with a 1.5% agarose pad. Representative images displayed. Arrows, live trophozoites; arrowheads, live pseudocysts.

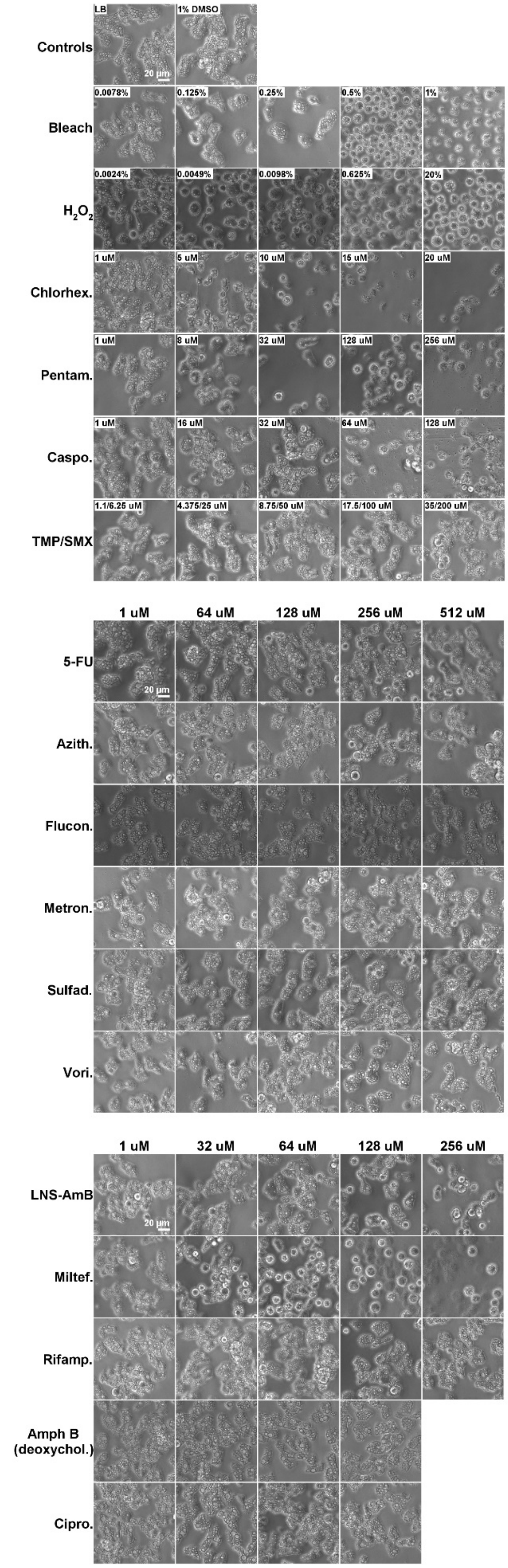

Controls
LB
1% DMSO
20 µm
Bleach
0.0078%
0.125%
0.25%
0.5%
1%
H2O2
0.0024%
0.0049%
0.0098%
0.625%
20%
Chlorhex.
1 uM
5 uM
10 uM
15 uM
20 uM
Pentam.
1 uM
8 uM
32 uM
128 uM
256 uM
Caspo.
1 uM
16 uM
32 uM
64 uM
128 uM
TMP/SMX
1.1/6.25 uM
4.375/25 uM
8.75/50 uM
17.5/100 uM
35/200 uM
1 uM
64 uM
128 uM
256 uM
512 uM
5-FU
20 µm
Azith.
Flucon.
Metron.
Sulfad.
Vori.
1 uM
32 uM
64 uM
128 uM
256 uM
LNS-AmB
20 µm
Miltef.
Rifamp.
Amph B
(deoxychol.)
Cipro.

**Figure S11: Morphological confirmation of cell phenotypes following drug treatment.** Microscopy of trophozoites in plate after 20-hour incubation with drugs or media, as described in figure 5. Phase images acquired before spent media was removed from wells using an EVOS FL Auto Imaging System (20x objective). Higher concentrations of caspofungin and pentamidine show a small amount of precipitated drug in the background of images. Selected representative images displayed—lowest and highest concentrations tested for all drugs, plus either the highest 3-4 concentrations or additional lower concentrations to show relevant morphological changes induced by active compounds.

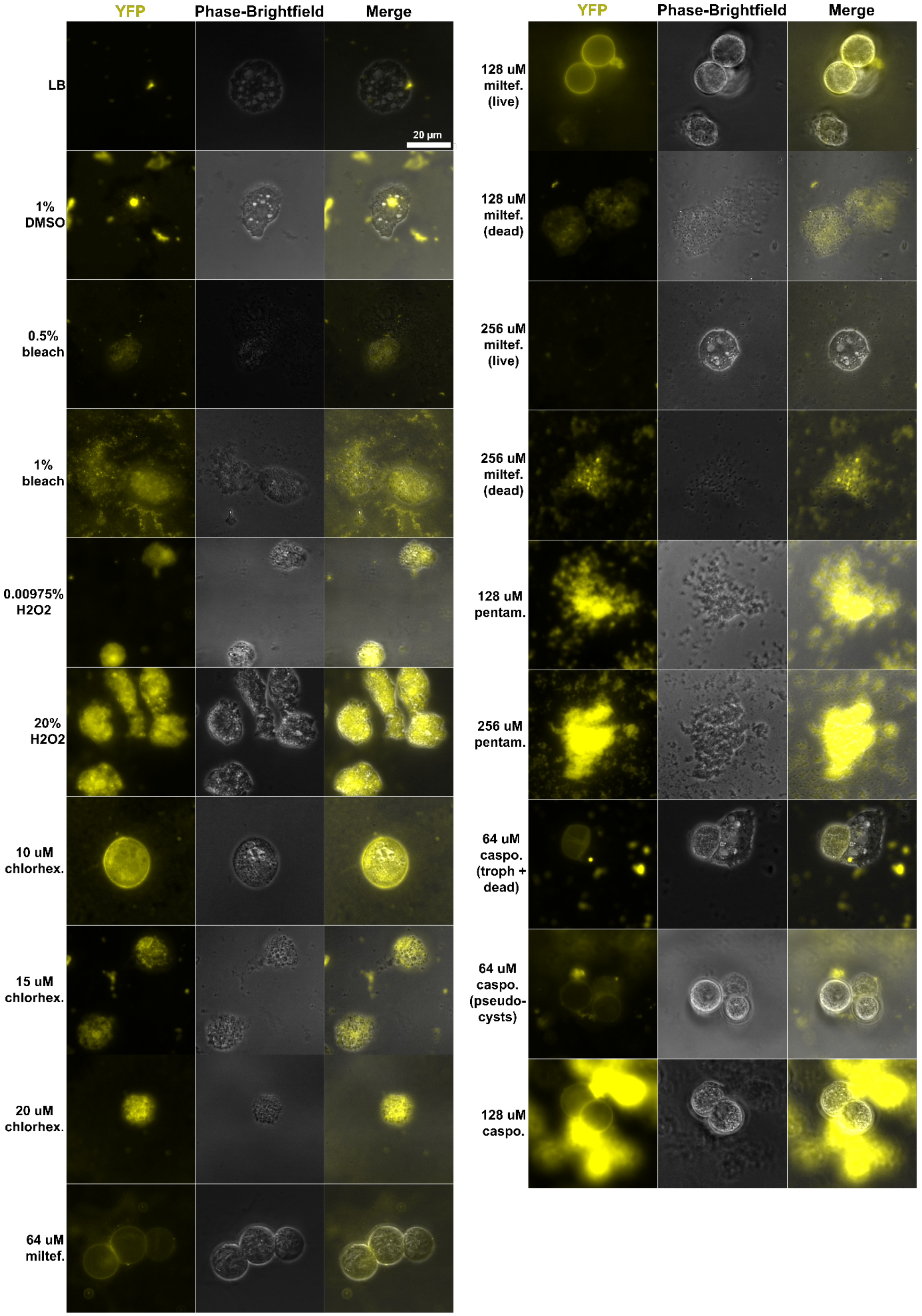
YFP
Phase-Brightfield
Merge
LB
20 µm
1% DMSO
0.5% bleach
1% bleach
0.00975% H2O2
20% H2O2
10 uM chlorhex.
15 uM chlorhex.
20 uM chlorhex.
64 uM miltef.
YFP
Phase-Brightfield
Merge
128 uM miltef. (live)
128 uM miltef. (dead)
256 uM miltef. (live)
256 uM miltef. (dead)
128 uM pentam.
256 uM pentam.
64 uM caspo. (troph + dead)
64 uM caspo. (pseudo-cysts)
128 uM caspo.

**Figure S12: Treatment with miltefosine and caspofungin results in pseudocyst formation.** $10^5$ trophozoites per well were incubated in a 96-well plate for 20 hours at 25°C in conditions as indicated. Spent media containing non-adherent cells was removed and transferred to an Eppendorf tube, stained with 100 µg/mL ConA-Alexa Fluor 488, and imaged using a Nikon Eclipse Ti-E inverted microscope (100x objective with oil) and a Hamamatsu ORCA-Fusion BT Digital CMOS camera in a MatTek glass-bottom dish overlaid with a 1.5% agarose pad. Selected drugs and concentrations were prepared as described in figure 5. Representative images displayed. The selected image for 10 µM chlorhexidine shows a dead pseudocyst, with granular appearance on phase contrast imaging and diffuse cytoplasmic staining with ConA.

### Supplemental methods

Media: Peptone Yeast Glucose medium (PYG): 2% (w/v) bacto peptone; 0.1% (w/v) bacto yeast extract; 0.1% (w/v) sodium citrate dihydrate;, 0.4 mM CaCl2; 4 mM MgSO4; 2.5 mM Na2HPO4; 2.5 mM KH2PO4; 0.05 mM Fe(NH4)2(SO4)2; 0.1 M glucose. 0.22 µm filter sterilized. Protocol available at https://dx.doi.org/10.17504/protocols.io.bvqbn5sn

Neff's first encystment medium(24) (EM): 0.1 M KCl; 0.02 M Tris base; 8 mM MgSO4; 0.4 mM CaCl2; 1 mM NaHCO3; adjusted to pH 8.5 with NaOH (Macron). Protocol available at https://dx.doi.org/10.17504/protocols.io.bvqcn5sw

Neff's modified second encystment medium(24) (EMb): 0.1 M KCl, 0.04 M NaHCO3, 8 mM MgSO4, 0.4 mM CaCl2, 0.32 mM 2-Amino-2-methyl-1,3-propanediol; pH 8-8.5 unadjusted.

PBS-MC: 17.4 mM $Na_2HPO_4$, 3.5 mM $NaH_2PO_4$ , 0.9 mM $CaCl_2$, 3.5 mM KCl, 0.9 mM $MgCl_2$, 0.137 M NaCl

Luria broth (LB): 10 g/L casein digest peptone, 10 g/L NaCl, 5 g/L yeast extract.

All media were sterilized by passage through a 0.22µm filter except for LB, which was autoclaved.
